# Glial scar formation by reactive astrocytes derived from oligodendrocyte progenitor cells after closed-head injury

**DOI:** 10.1101/2024.09.16.613178

**Authors:** Hinata Matsuda, Erika Tsuji, Agnes Antonetta Orchidia Riberu, Hideyuki Okano, Mitsuhiro Morita

## Abstract

The diversity of reactive astrocytes is key to understanding complicated pathological processes in the brain. The accumulation of reactive astrocytes expressing the neural stem/precursor cell marker Nestin is common after brain injury, but the pathological implications of this reactive astrocyte subpopulation remain elusive. This study initially aimed to determine the origin and fate of these reactive astrocytes expressing Nestin by characterizing cells labeled with green fluorescent protein (GFP) after closed-head injury, using a *Nestin* promoter region widely utilized to study neural stem/precursor cells. Unexpectedly, oligodendrocyte progenitor cells (OPCs), rather than astrocytes, were robustly and selectively labeled with GFP. A fraction of these cells showed a subsequent upregulation of astrocyte markers and were incorporated into glial scars. These glial scars are aggregates of reactive astrocytes that form between lesion cores and the perilesional recovering region. Deletion of the *Stat3* gene, which is essential for astrocyte activation, using a *Nestin* promoter reduced glial scars, further confirming that OPCs are involved in glial scar formation. Reactive astrocytes labeled with a *glial fibrillary acidic protein* promoter differed in morphology and distribution from astrocytes derived from OPCs. This confirms that astrocytes and OPCs produce distinct reactive astrocyte subpopulations. Some GFP-labeled OPCs lacking astrocyte markers were found to distribute in perilesional recovering regions. The reduced expression of Nestin and OPC markers in these non-astrocytic descendants of OPCs, coupled with a significant fraction of these cells remaining olig2-positive, suggests that OPCs give rise to both reactive astrocytes and oligodendrocytes. These findings suggest that OPCs are activated by a novel process after brain injury.

## 1. Introduction

Hypertrophy, proliferation and metabolic changes in astrocytes, a condition called reactive astrocytosis, are hallmarks of neurological disorders and brain injuries (Sofroniew, 2009). Our previous studies suggest the diversity of activation processes and pathological implications of reactive astrocytes (Morita et al., 2003; Sawant et al., 2024; Suzuki et al., 2012). The accumulation of reactive astrocytes can be beneficial or detrimental, depending on the disease type and progression (Anderson et al., 2014). A transcriptome analysis of inflammation and stroke model mice identified neurotoxic and neuroprotective subpopulations of reactive astrocytes (Zamanian et al., 2012), which were later named A1 and A2 reactive astrocytes, respectively (Liddelow et al., 2017). Furthermore, the advance of spatial and temporal resolution in transcriptomics by single-cell RNA sequencing is revealing the remarkable heterogeneity of reactive astrocytes among neurological diseases (Matusova et al., 2023). Because optimizing the density and function of reactive astrocytes likely leads to improved outcomes in patients with brain disorders, the origins, functional molecules, and cell signaling of reactive astrocytes are under intense investigation as potential diagnostic and/or therapeutic targets (Finsterwald et al., 2015).

To investigate astrocyte responses to localized brain tissue degeneration, we have developed a novel closed-head injury model and named the “photo injury” mouse (Suzuki et al., 2012). Because focal cerebral lesions were generated by illumination with intense light through a thinned-skull cranial window, this model avoids artificial inflammation due to craniotomy, a procedure commonly used in conventional models of traumatic brain injury (Peterson et al., 2015) and induces gliosis by itself (Xu et al., 2007). Furthermore, the degenerative process after photo injury has been attributed to localized lesions, but not to long-lasting metabolic changes common in the widespread ischemic penumbras of stroke models (Strong et al., 1988). Thus, this model allows the study of astrocyte reactivities associated exclusively with tissue degeneration and recovery. An early study identified two distinct subpopulations of reactive astrocytes after photo injury (Suzuki et al., 2012): proliferating and Nestin-expressing reactive astrocytes in perilesional recovering regions and non-proliferating and Nestin-negative hypertrophic reactive astrocytes in distal regions. In addition, the ablation of perilesional Nestin-expressing reactive astrocytes accelerates blood-brain barrier (BBB) disruption and increases inflammation, leading to impaired wound healing (Sawant et al., 2024). Thus, the essential anti-inflammatory and regenerative functions of the perilesional Nestin-expressing reactive astrocytes were suggested.

This study was initially designed to address a hypothesis that the expression of the neural stem/precursor cell marker, Nestin, is associated with de-differentiation or trans-differentiation of reactive astrocytes. For this purpose, the origin and fate of perilesional Nestin-expressing reactive astrocytes were analyzed by labeling or manipulating these cells by using a *Nestin* promoter. As a result, we unexpectedly found that NG2-positive oligodendrocyte progenitor cells (OPCs) at the lesion core were robustly and exclusively labeled by the *Nestin* promoter, and subsequently produced reactive astrocytes. Furthermore, the morphology, distribution, and putative function of reactive astrocytes derived from OPCs were distinct from those derived from astrocytes, which were labeled by a *glial fibrillary acidic protein* (*Gfap*) promoter. The present study demonstrates a novel pathway for generating reactive astrocytes in the perilesional region after closed head injury and proposes a unique therapeutic and diagnostic target for improving the outcome of brain injury.

## 2. Materials and methods

### 2.1 Animal experiments

All animal experiments were approved by the Institutional Animal Care and Use Committee of Kobe University (Permission number: 28-06-06) and were performed according to the Animal Experimentation Regulations of Kobe University. *Nestin-CreER^T2^;Rosa26-EGFP* and *Nestin-CreER^T2^;Stat3^f/f^;Rosa26-EGFP* mice were generated by crossing *Nestin-CreER^T2^* mice (Imayoshi et al., 2006a) with *Rosa26-EGFP* mice (Kawamoto et al., 2000) and *Stat3*^f/f^ mice (Okada et al., 2006), respectively, whereas *Gfap-CreER^T2^/Rosa26-GCaMP6* mice were generated by crossing *Gfap-CreER^T2^* mice (Hirrlinger et al., 2006) and *Rosa-GCaMP6* mice (Ohkura et al., 2012). Photo injury was generated in 6–10-week-old mice of both sexes as described (Suzuki et al., 2012). Briefly, a thinned-skull cranial window (∼ 0.5 mm in diameter and ∼ 20 μm in thickness) was created over the right primary somatosensory cortex (1.5 mm posterior to the bregma and 3 mm lateral from the midline) by scraping the skull using micro-drill burrs (19007-07M; FST, Foster City, CA, USA) and a microsurgical blade (Nordland Blade #6900; Salivan, Charlotte, NC, USA). The mouse was mounted on a microscope stage for light illumination. Its head was fixed with the cranial window centered under a 20× water immersion microscope objective (XLUMPLFLN 20× W NA1.0; Olympus, Tokyo, Japan), setting the focal point 200 μm below the cortical surface. The cortical tissue was exposed for a duration of 1.5–2.5 min to the light from a 90-W halogen bulb mounted in a housing directly over a microscope camera port. Tamoxifen (Toronto Research Chemicals, Toronto, Canada) dissolved in corn oil (Sigma-Aldrich, St. Louis, MO, USA) was orally administered (250 mg/kg). For bromodeoxyuridine (BrdU) labeling, BrdU (50 mg/kg, i.p.) was administered on days 1 ∼ 6. All mice were housed under a 12-h light: 12-h dark cycle with food and water available ad libitum.

### 2.2 Immunofluorescence analysis

Mice were sacrificed at given time points, and tissue samples were subjected to histological analysis. After anesthesia with a sodium pentobarbital overdose (75 mg/ kg, i.p), the mice were transcardially perfused with ice-cold 0.1 M PBS containing 1000 units/L heparin (Wako, Tokyo, Japan), followed by 4% (w/v) paraformaldehyde in 0.1 M phosphate-buffered saline (PBS). Brains were post-fixed for six hours, and coronally sliced at a thickness of 100 μm in PBS using a microslicer (Zero-One, DOSAKA, Osaka, Japan). The slices were stored at −20°C in a preservative containing 0.3% glycerol, 0.3% ethylene glycol, 0.1 M sodium phosphate and, 0.85% NaCl. For immunofluorescence staining, the slices were washed twice with PBS for 10 min each, treated with 50 mM NH_4_Cl for 10 min, washed twice with PBS for 10 min each, permeabilized with PBS containing 0.4% (w/v) Triton X-100 for two hours, and blocked with a blocking buffer (PBS containing 5% normal goat serum and 0.1% Triton X-100). The sections were incubated overnight at 4°C with primary antibodies in blocking buffer, returned to room temperature, washed three times with PBS, incubated with secondary antibody in blocking buffer for two hours, washed three times with PBS for 10 min each, and mounted with gelvatol (Harlow and Lane, 2006) containing 100 mg/ml DABCO (Sigma).

Primary antibodies included guinea pig polyclonal anti-GFAP (1:400; a gift from Prof. Seiji Miyata, Kyoto Institute of Technology); rabbit anti-S100β (1:400; DAKO, Glostrup, Denmark), chicken anti-Nestin (1:1000; Novus Biologicals, Centennial, CO. USA), rabbit andti-NG2 (1:200; Chemicon, Temecula, CA, USA), rabbit anti-Sox2 (1:100; Millipore, Temecula, CA, USA), rabbit anti-PDGFRα (1:100; Santa Cruz Biotechnology, Dallas, TX, USA), goat anti-PDGFRα (1:40; R&D Systems, Minneapolis, MN, USA), goat anti-PDGFRβ (1:200; R&D Systems), rabbit anti-Olig2 (1:250; Abcam), rabbit anti-Iba1 (1:200; Wako), and rat anti-BrdU (1:100; Accurate Chemical, Westbury, NY, USA). Immunofluorescence was visualized using FITC-, Cy3-or Cy5-conjugated secondary antibodies (1:200; Jackson ImmunoResearch Laboratories, West Grove, PA, USA). For BrdU double staining, samples were treated with 2 M hydrochloric acid (37 °C, 30 min), and then neutralized with 0.1 M sodium borate (pH 8.5, 10 min). Z-stack confocal fluorescence images (step size = 2 μm) were obtained by Fluoview1000 (Olympus, Tokyo, Japan) and analyzed by Fiji (Schindelin et al., 2012).

### 2.3 Quantitative analysis

For quantitative analysis, slices showing the maximum horizontal distance of the lesion, presumably including its central region, from three or four mice were subjected to immunofluorescence staining, with one image from each slice measured and included in subsequent statistical analysis by Excel or SciPy library in Python.. To determine the ratio of cell type marker or BrdU positive cells to GFP positive cells, 20 randomly selected GFP positive cells were analyzed.

### 2.4 Chemicals

Unless otherwise indicated, all chemicals were obtained from Nacalai tesque (Kyoto, Japan).

## 3. Results

### 3.1 GFP-labeling of non-astrocytic cells and scar-forming reactive astrocytes in *Nestin-CreER^T2^:Rosa26-GFP* mice after brain injury

*Nestin-CreER^T2^;Rosa26-GFP* mice subjected to photo injury were treated with tamoxifen one day post-injury (1 dpi), and the distributions of GFP-labeled cells and reactive astrocytes were examined by immunofluorescence staining of GFP, GFAP and S100β. As shown in low magnification images of lesions (Fig. 1A), the majority of GFP+ cells were within the inner region devoid of astrocytes (lesion core, LC) at 4 dpi, indicating the induction of GFP expression in non-astrocytic cells in the LC during the early stage of injury. In contrast, GFP+ cells were in the perilesional region containing astrocytes at 14 dpi. As described in our previous study (Suzuki et al., 2012), the reduction of LC size between 4 dpi and 14 dpi reflects the migration of perilesional reactive astrocytes into the LC. Thus, the co-distribution of GFP+ cells and astrocytes in the later stage is likely due to the migration of reactive astrocytes into the area where GFP+ cells were generated. GFP+ cells were further analyzed with higher magnification images at 4, 7, 14, and 28 dpi (Fig. 1B). S100β+ normal astrocytes were found in the contralateral hemisphere at 28 dpi, whereas GFAP+ reactive astrocytes and GFP-labeled cells were not, confirming the lack of astrocyte activation or nestin expressing cells in normal regions. At 4 dpi, ipsilateral GFP+ cells were GFAP-/S100β-(empty arrowheads) and localized to the area surrounded by perilesional GFAP+/S100β+ reactive astrocytes. We defined the region covered by reactive astrocyte processes that extend radially from the LC (arrows) after 7 dpi as the perilesional region (PR), because these process-bearing reactive astrocytes were shown to be Nestin+ in our previous study (Suzuki et al., 2012). GFP+ cells were found not only in the LC, but also in the PR, with most of these cells remaining GFAP-/S100β-at 7 dpi. In addition, GFP+ cells in the PR showed a uniform spherical morphology. The appearance of GFP+ cells in the PR most likely reflects the reduction in volume of the LC accompanied by the migration or process extension of reactive astrocytes into the LC (Suzuki et al., 2012). Aggregates of reactive astrocytes lacking radial processes formed at the interface between the LC and RP at 14 dpi, and they included GFP+/GFAP+/S100β+ cells (filled arrowheads). These aggregates are designated as glial scars (GS), which are glial structures at interfaces between neuronal and non-neuronal tissues. This designation is supported by our previous study, which showed high numbers of neurons in the PR and a complete loss of neurons in the LC (Suzuki et al., 2012). The appearance of aggregates was accompanied by the disappearance of GFP+ cells from the LC, suggesting the conversion of the GFP+ cells to scar-forming reactive astrocytes. The glial scars composed of GFP+/GFAP+/S100β+ cells, whereas the morphology and immunoreactivity of GFP+(/GFAP-/S100β-) cells in the PR remained the same at 28 dpi. These results indicate that a fraction of cells labeled with GFP by the *Nestin* promoter becomes reactive astrocytes forming glial scars, with the remainder remaining as non-astrocytic cells in the PR.

**Figure 1.**
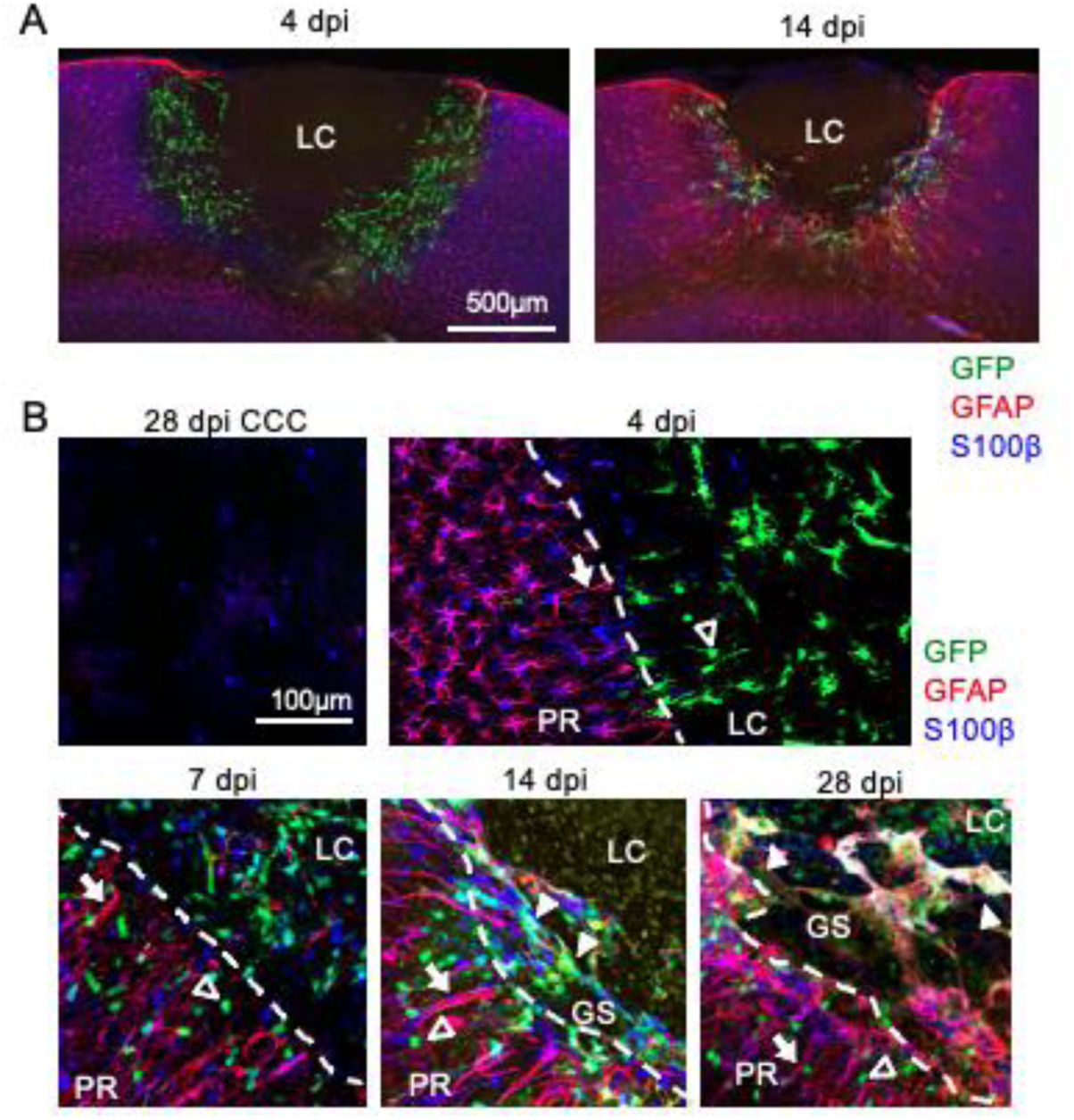
GFP labeling of non-astrocytic cells and scar-forming reactive astrocytes in *Nestin-CreER^T2^:Rosa26-GFP* mice after brain injury. *Nestin-CreER^T2^:Rosa26-EGFP* mice were subjected to photo injury and treated with tamoxifen at 1 dpi, followed by immunofluorescence analysis of expression of GFP, S100 and GFAP. Low magnification images of lesion core (LC) at 4 and 14 dpi (A). High magnification images of astrocytes and GFP+ cells in the contralateral cortex at 28 dpi and in the ipsilateral cortex at 4, 7, 14 and 28 dpi (B). The borders between the LC and PR are indicated by dashed lines, and the accumulation of GFP+ cells expressing astrocytic markers at the interface between LC and PR was defined as glial scar (GS). Representative reactive astrocytes extending fibers radially from the LC (arrows), GFP+ cells lacking astrocyte markers (empty arrowheads) and GFP+ scar-forming reactive astrocytes (filled arrowheads) are shown.

### 3.2 Generation of scar-froming reactive astrocytes from non-astrocytic cells

GFP+ reactive astrocytes in the GS are likely derived from the non-astrocytic GFP+ cells that appeared earlier. However, administration of tamoxifen at 1 dpi may have induced GFP labeling in reactive astrocytes between 7 and 14 dpi. To address the timing of GFP labeling, *Nestin-CreER^T2^:Rosa26-GFP* mice were treated with tamoxifen at 1, 4, or 7 dpi, and GFP labeling was examined by immunofluorescence staining of GFP and GFAP at 14 dpi (Fig. 2). Few GFP+ cells were found in the mice treated with tamoxifen at 4 and 7 dpi (Fig. 2A), with the percentages of GFP+ cells induced by tamoxifen at 4 dpi and 7 dpi being 19.7 ± 3.0% and 14.6 ± 9.1%, respectively, of these cells at 1 dpi, indicating that tamoxifen treatment after 4 dpi significantly reduces GFP labeling (Fig. 2B). The number of GFP+ cells induced by tamoxifen at 1dpi may be underestimated, and the percentages at 4 dpi and 7 dpi be even lower, because individual GFP+ cells in astrocyte aggregates in the GS were sometimes difficult to count separately at 1 dpi, while sparse and easy to count at 4 dpi and 7 dpi. Collectively, tamoxifen treatment at 1 dpi induces GFP labeling between 1 and 4 dpi, confirming that the GFP+ cells in Figure 1 reflect the fate of non-astrocytic GFP+ cells appearing at 4 dpi.

**Figure 2.**
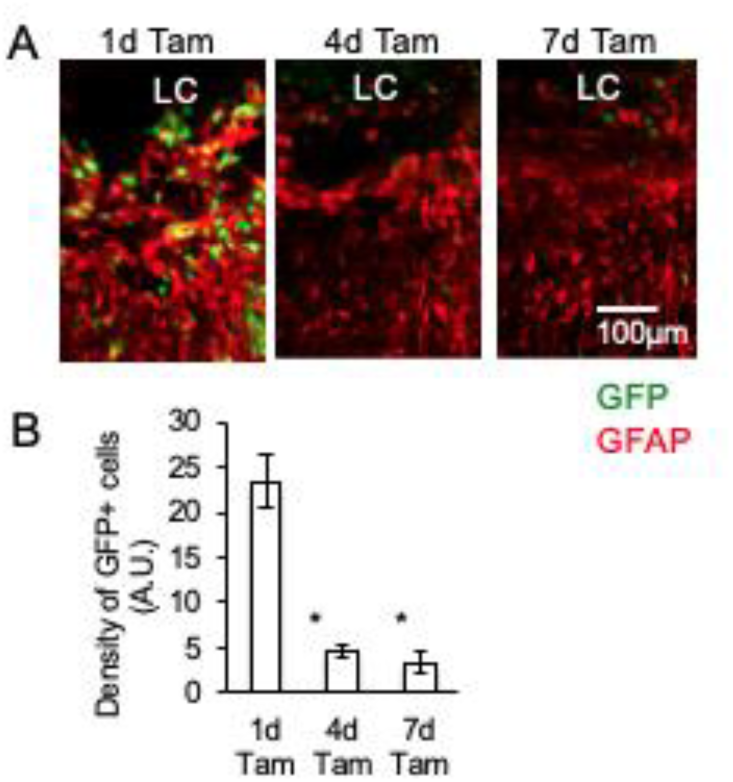
Tamoxifen treatment induced GFP labeling exclusively during early stages after brain injury. *Nestin-CreERT2:ROSA26-EGFP* mice were subjected to photo injury and treatment with tamoxifen at 1 dpi (1d Tam), 4 dpi (4d Tam) and 7 dpi (7d Tam), followed by immunofluorescence analysis of GFP and GFAP at 14 dpi. A. Representative images of lesions. B. Density of GFP+ cells. Mean ± SD, n = 3 images from three mice for each value, *p<0.05 by ANOVA and Dunnett’s multiple compared with results at 1d Tam.

### 3.3 GFP-labeling of oligodendrocyte precursor cells (OPCs) in in *Nestin-CreER^T2^:Rosa26-GFP* mice after brain injury

The origin of GFP+ cells was examined by immunofluorescence staining of GFP and cell type markers at 4 dpi (Fig. 3). Various cell types in the LC and PR were found to be Nestin+, consistent with studies reporting Nestin expression in reactive astrocytes (Suzuki et al., 2012), as well as in pericytes (Nakata et al., 2017) and oligodendrocyte precursor cells (OPCs) (Liu et al., 2015) (Fig. 3A). The widespread Nestin immunoreactivity in the LC at 4 dpi was not mentioned in our previous study (Suzuki et al., 2012), because that study focused on the PR during later stages (> 7 dpi). Most GFP+ cells were Nestin+, whereas most perilesional Nestin+ cells were morphologically similar to reactive astrocytes at 4 dpi and lacked GFP labeling. This lack of GFP labeling indicates that the *Nestin* promoter region used in *Nestin*-CreER^T2^ mice, consisting of 5.8 kb of the 5’ flanking region and 1.8 kb including the second intron of rat *Nestin* gene (Imayoshi et al., 2006b), is not active in all cell types expressing Nestin after brain injury. Because this promoter region including an essential second intrinsic enhancer is active in both embryonic and adult neural stem/progenitor cells *in vivo* (Lagace et al., 2007; Zimmerman et al., 1994), GFP labeling may be associated with dedifferentiation to neural stem cells. Another neural stem cell marker, Sox2, was also found to be expressed in both GFP+ and perilesional GFP-cells (Fig. 3B), indicating that the *Nestin* promoter induced GFP labeling in a subset of cells positive for neural stem/progenitor cell marker. The immunoreactivities of astrocyte markers GFAP and S100β localized in the PR, whereas and GFP+ cells were GFAP-/S100β-, confirming that these GFP+ cells were not derived from reactive astrocytes (Fig. 3C and D). The immunoreactivity of neuron-glial antigen 2 (NG2), a marker of OPCs and pericytes (McTigue et al., 2006; Nishiyama et al., 1996), was also assessed, because these cell types show upregulation of Nestin expression (Liu et al., 2015; Nakata et al., 2017). Various cell types in the LC, including GFP+ cells, were found to be NG2+, suggesting that GFP+ cells were pericytes or OPCs (Fig. 3E). Assessment of immunoreactivity for PDGFRα and PDGFRβ, which are markers for OPCs and pericytes, respectively (Fig. 3F and G), showed that GFP+ cells were PDGFRα+/PDGFRβ-, indicating that they were OPCs. This finding was further confirmed by the nuclear localization of Olig2, a marker of developing OPCs, in GFP+ cells (Fig. 3H). Olig2+ was also expressed in perilesional cells, presumably reflecting its expression in proliferating reactive astrocytes (Chen et al., 2008). Quantitative analysis of these immunofluorescence data indicated that the majority of GFP+ cells were Nestin+ (95.0 ± 4.1%), NG2+ (80.0 ± 4.1%) and PDGFRα+ (96.7 ± 4.7%), while significantly fewer GFP+ cells were GFAP+ (5.0 ± 4.1%), S100β-(26.7 ± 9.4%) and PDGFRα+ (16.7 ± 17.0%), further confirming that GFP+ cells were derived from OPCs.

**Figure 3.**
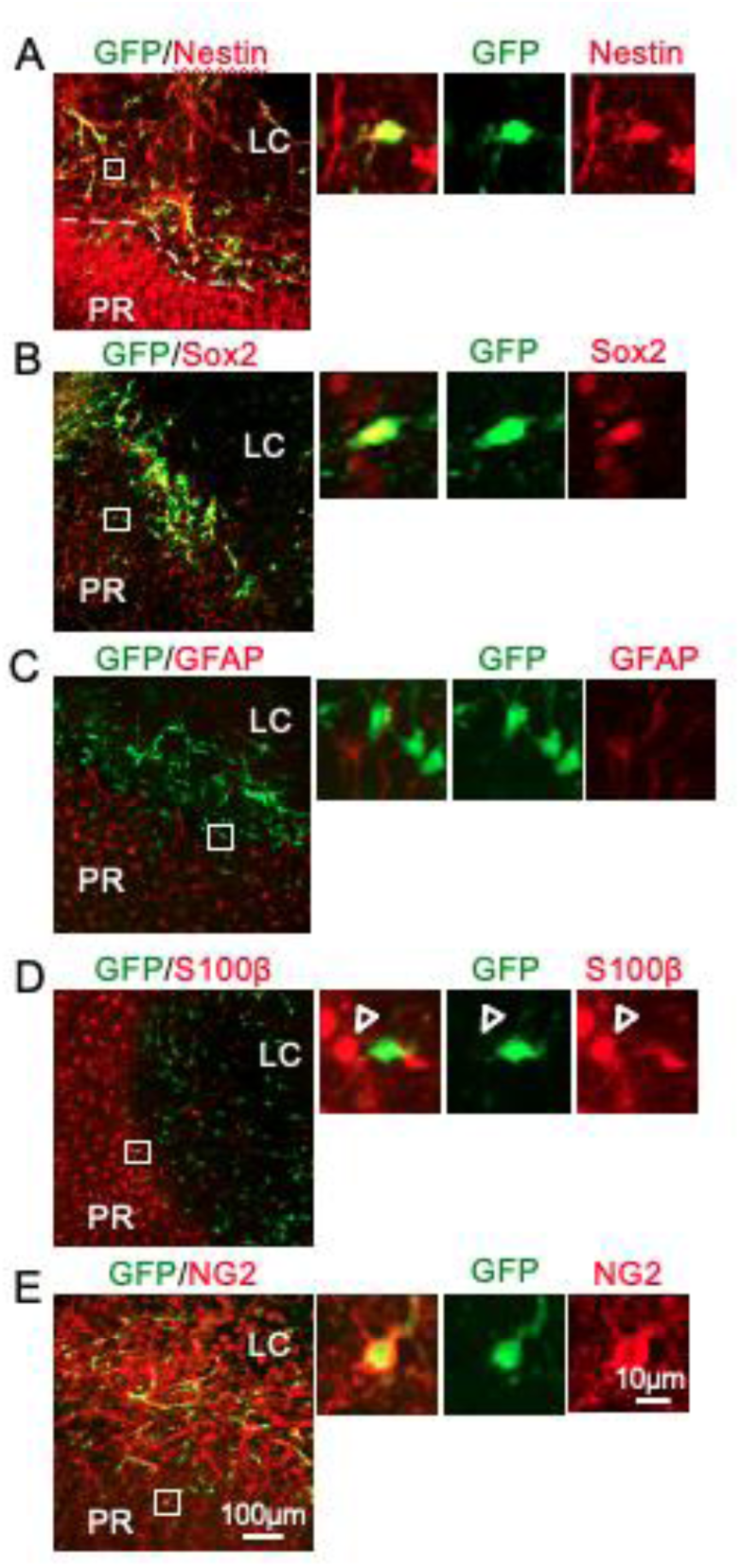
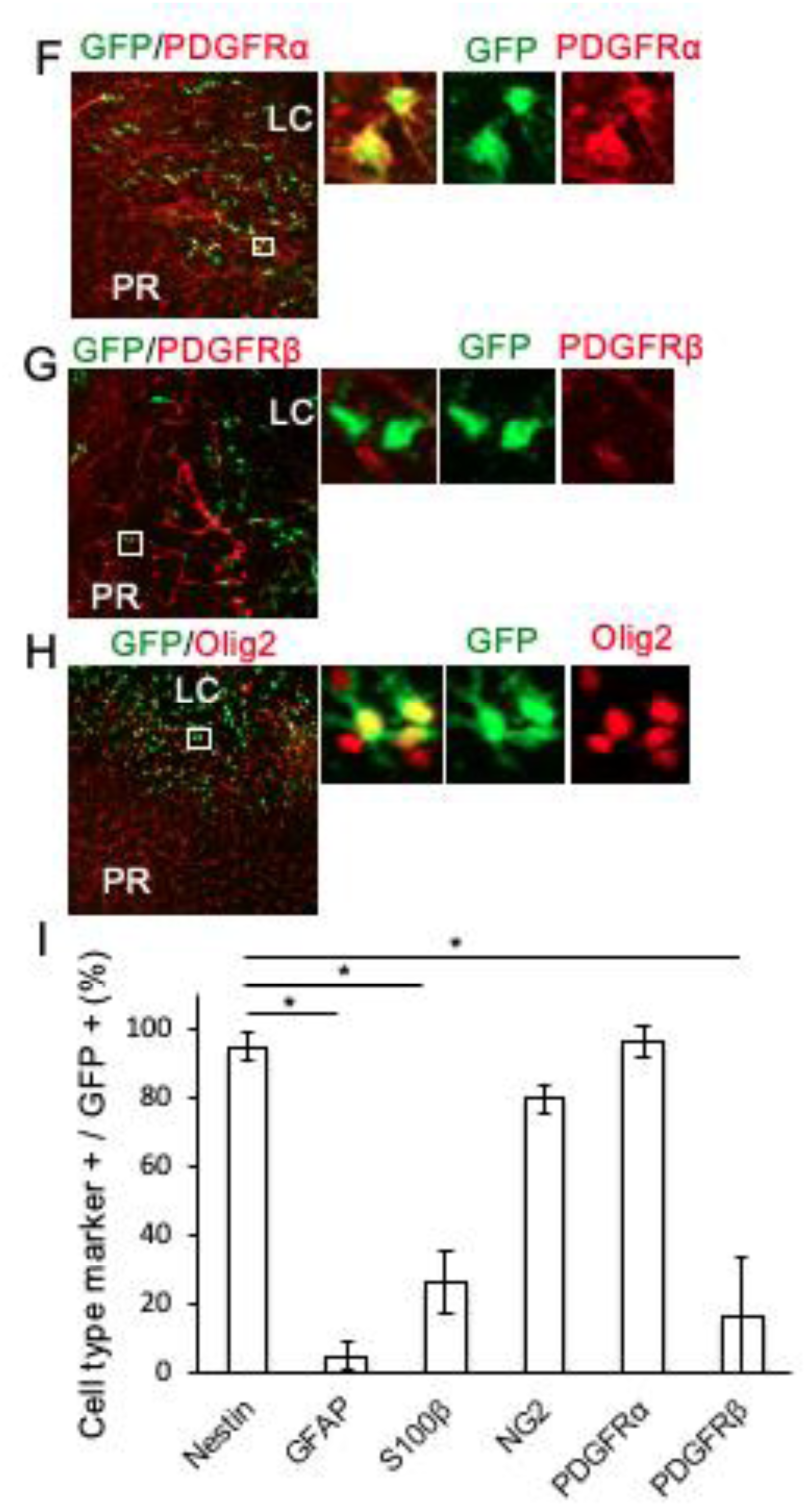
Derivation of GFP+ cells from OPCs. *Nestin-CreER^T2^:Rosa26-EGFP* mice were subjected to photo injury and treatment with tamoxifen at 1 dpi, followed by immunofluorescence analysis of GFP and cell type markers at 4 dpi. Representative images of GFP and Nestin (A), Sox2 (B), GFAP (C), S100β (D), NG2 (E), PDGFR (F), PDGFR (G) and Olig2 (H). Representative GFP+ cells highlighted by white boxes in the low magnification images on the left hand side were enlarged in the three images on the right side. I. Ratios of cell type marker positive cells to GFP+ cells. Mean ± SD, n = 60 GFP+ cells from three mice. *P < 0.05 compared with Nestin by ANOVA and Dunnett’s multiple comparison test.

### 3.4 Reductions of GFP+ cells and glial scar formation by conditional *Stat3* knockout

The *Stat3* gene is essential for both the differentiation and pathological activation of astrocytes (Cattaneo et al., 1999; Herrmann et al., 2008). Thus, conditional knockout of *Stat3* using *Nestin*-*CreER^T2^:Stat3^f/f^:Rosa26-EGFP* mice should reduce the conversion of GFP+ cells to scar-forming reactive astrocytes. To test this hypothesis, *Nestin-CreER^T2^;Rosa26-GFP* (Cont) and *Nestin-CreER^T2^:Stat3 ^f/f^:Rosa26-EGFP* (STAT3 KO) mice were subjected to photo injury and treated with tamoxifen at 1 dpi, followed by immunofluorescence analysis of GFP and GFAP at 4, 14 and 28 dpi (Fig. 4). The numbers of GFP+ cells were markedly reduced at all time points (Fig. 4A). The density of GFP+ cells at 4 dpi was significantly lower in STAT3 KO (2.0 ± 0.1 A.U.) than in control (4.6 ± 0.8 A.U.) mice (Fig. 4B). Because GFP+ cells in the GS formed aggregates and were difficult to count after 14 dpi, the number of non-astrocytic GFP+ cells in the PR and the area occupied by astrocytic GFP+ cells in the GS were quantified separately at later time points. The reduction of perilesional GFP+ cells was quantitively confirmed until 28 dpi, as the density of perilesional GFP+/GFAP-cells was significantly lower in STAT3 KO (0.8 ± 0.5 A.U.) than in control (2.6 ± 0.3 A.U.) mice at 28 dpi (Fig. 4C). The involvement of the *Stat3* gene in scar formation was examined by comparing the areas of the GS (GFAP+ aggregates between the PR and the LC) normalized to the length of the corresponding LC border in control and STAT3 KO mice. The GS at 14 dpi were significantly smaller in STAT3 KO (15.5 ± 5.1 A.U) than in control (36.1 ± 5.4 A.U) mice (Fig. 4D). Scar formation at 28 dpi was not analyzed quantitatively, because the LC and GS structures were easily damaged during the immunofluorescence staining of these slices, presumably reflecting the degradation of these lesions. The area occupied by astrocytic GFP+/GFAP+ cells in the GS was significantly smaller in STAT3 KO (11.2 ± 7.0 %) than in control (49.7 ± 5.7 %) mice at 14 dpi (Fig. 4E), indicating the GS of STAT3 KO includes less GFP+ cells. Taking together, these results showed that *Stat3* knockout attenuated the accumulation of GFP+ cells during the initial stage after brain injury (< 4 dpi), reduced GFP+/GFAP-cells in the PR and GFP+/GFAP+ cells in glial scars, and inhibited glial scar formation.

**Figure 4.**
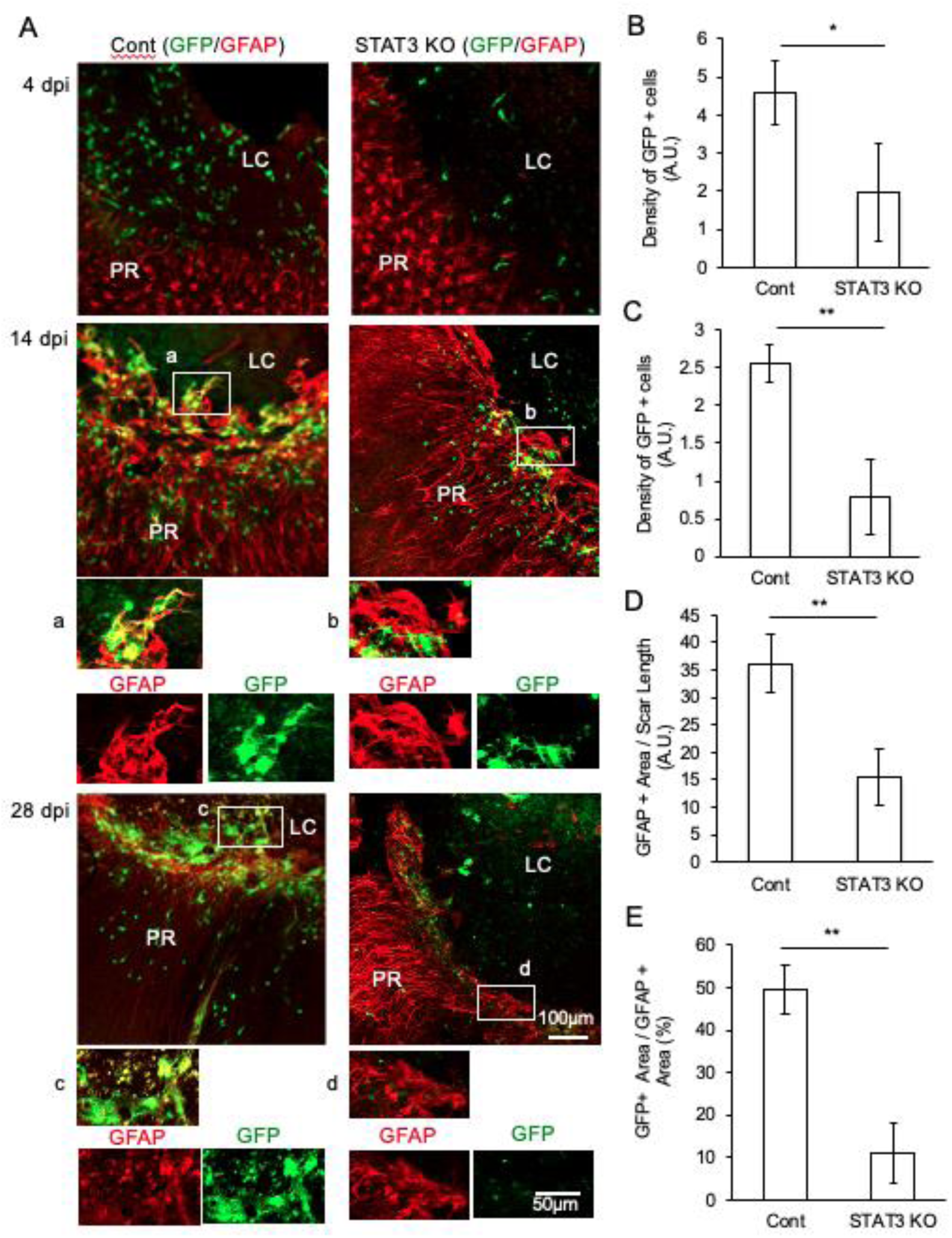
*Stat3* knockout reduced GFP+ cells and glial scar formation. *Nestin-CreERT2;Rosa26-GFP* (Cont) and *Nestin-CreERT2:Stat3^f/f^:ROSA26-EGFP* (STAT3 KO) mice were subjected to photo injury and treatment with tamoxifen at 1 dpi, followed by immunofluorescence analysis of GFP and GFAP at 4, 14 and 28 dpi. A. Representative immunofluorescence images. Scar-forming reactive astrocytes highlighted in the white boxes in the low magnification images at 14 and 28 dpi (a-d) were enlarged below. B. Density of GFP+/GFAP-cells at 4 dpi. Mean ± SD, n = 4 mice, *P < 0.05 by Student’s t-test. C. Density of GFP+/GFAP-cells in the PR at 28 dpi. n = 3 mice, **P < 0.01. D. Glial scar areas, defined as GFAP+ areas in the GS normalized by the length of the corresponding border of the LC, at 14 dpi. n = 3 or 4 mice, **P < 0.01. E. Proportions of areas occupied by cells labeled with GFP (GFP+ areas) in glial scars (GFAP+ areas). n = 3 or 4 mice, **P < 0.01.

### 3.5 Suppression of GFP+ cell proliferation by conditional Stat3 knockout

The robust reduction of GFP+ cells caused by *Stat3* knockout became prominent in the early stages of brain injury (4 dpi) and persisted until 28 dpi. As GFP-labeling and *Stat3* knockout by *Nestin-CreER^T2^*undergoes simultaneously after tamoxifen treatment, *Stat3* knockout less likely affects the number of cells, which induced GFP expressions. Thus, *Stat3* knockout may have suppressed the proliferation of GFP+ cells before 4dpi. To test this hypothesis, the proliferation of GFP+ cells in control and STAT3 KO mice was tested using BrdU (Fig. 5). GFP+/BrdU+ and GFP+/BrdU-cells were found in both control and STAT3 KO mice (Fig. 5A), although the proportion of GFP+ cells that were also BrdU+ was significantly lower in STAT3 KO (45.0 ± 7.1%) than in control (63.3 ± 4.7%) mice (Fig. 5B), indicating that OPC proliferation is STAT3-dependent following *Nestin* gene activation by brain injury.

**Figure 5.**
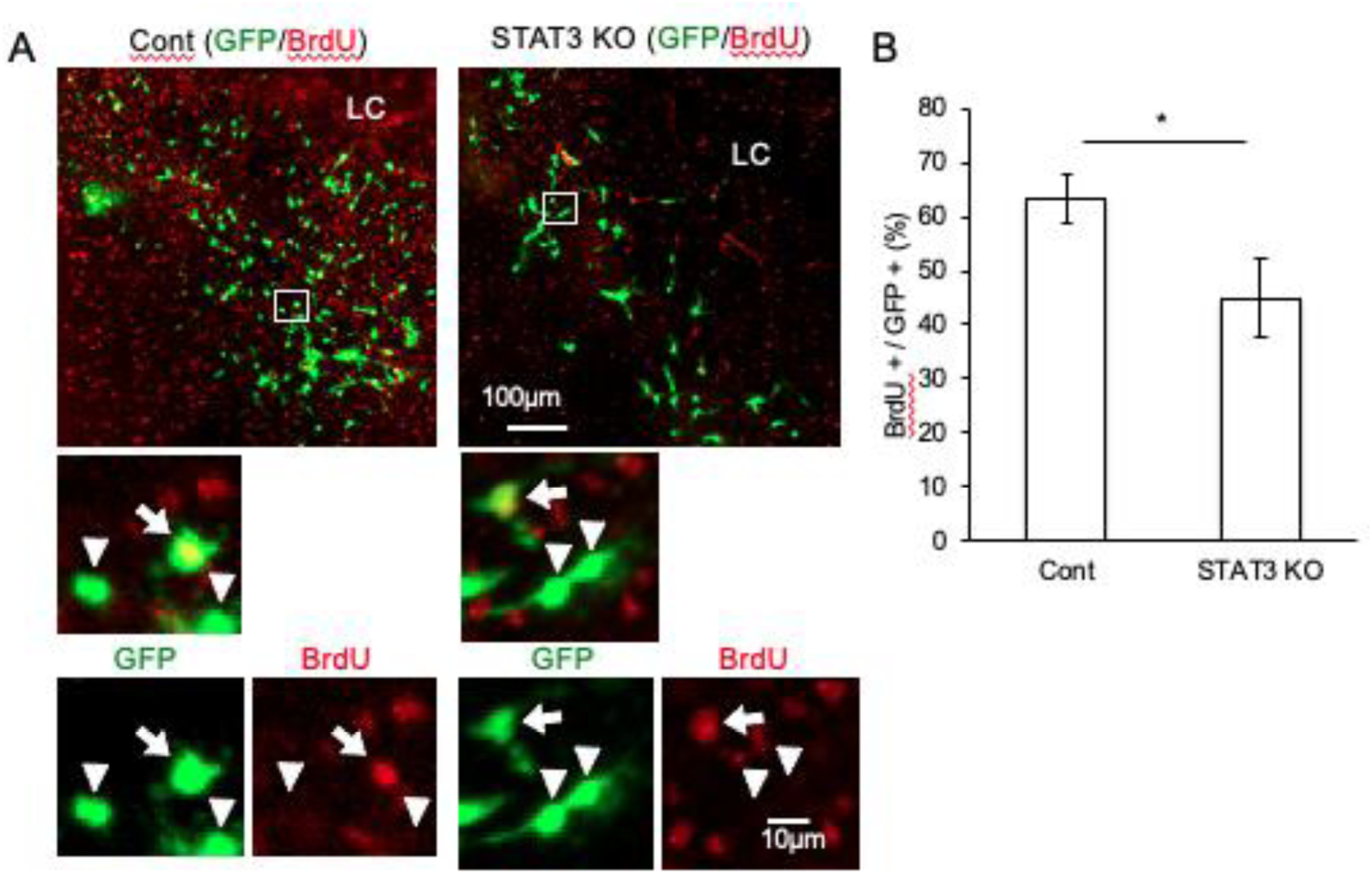
*Stat3* knockout suppressed the proliferation of GFP+ cells. Control and STAT3 KO mice were subjected to photo injury, administered tamoxifen at 1 dpi and intraperitoneal BrdU (50 mg/kg) at 1 ∼ 3 dpi, followed by immunofluorescence staining for GFP and BrdU at 4 dpi. A. Representative immunofluorescence images. GFP+ cells highlighted by the white boxes in low magnification images were enlarged below. GFP+/BrdU+ cells and GFP+/BrdU-cells are indicated by arrows and arrowheads, respectively. B. Ratios of GFP+/BrdU+ cells to total GFP+ cells. Mean ± SD, n = 60 cells from 3 mice, *P < 0.05 by Student’s t-test.

### 3.6 Lack of astrocyte-derived reactive astrocytes in glial scar

The finding that GFP+ reactive astrocytes in the GS were morphologically distinct from GFAP+ reactive astrocytes in the PR suggested that OPCs and astrocytes give rise to different reactive astrocyte subpopulations. This hypothesis was tested by using *Gfap-CreERT2/ROSA26-GCaMP6* (GFAPp) mice, which allow the detection of astrocyte-derived reactive astrocytes by GFP immunostaining. GFAPp mice were subjected to photo injury and treated with tamoxifen at 1 dpi, followed by immunofluorescence analysis for GCaMP6 and GFAP at 14 dpi. GCaMP6+ cells were found to be distributed in the PR as reactive astrocyte extending processes from the LC, equivalent to the Nestin+ reactive astrocytes in our previous study (Suzuki et al., 2012) (Fig. 6A upper and Aa). In contrast, few GCaMP6+ cells were found in glial scars (Fig. 6Ab). Quantitative comparisons showed that significantly smaller areas of the GS were occupied by GFP+ cells in GFAPp (3.9 ± 2.0%) than control (*Nestin-CreER^T2^: Rosa26-EGFP*, 43.7 ± 6.9%), indicating *Gfap* promoter is less potent in labeling reactive astrocyte in the GS than *Nestin* promoter. Thus, scar-forming reactive astrocytes did not derive from astrocytes, but were generated from OPCs.

**Figure 6.**
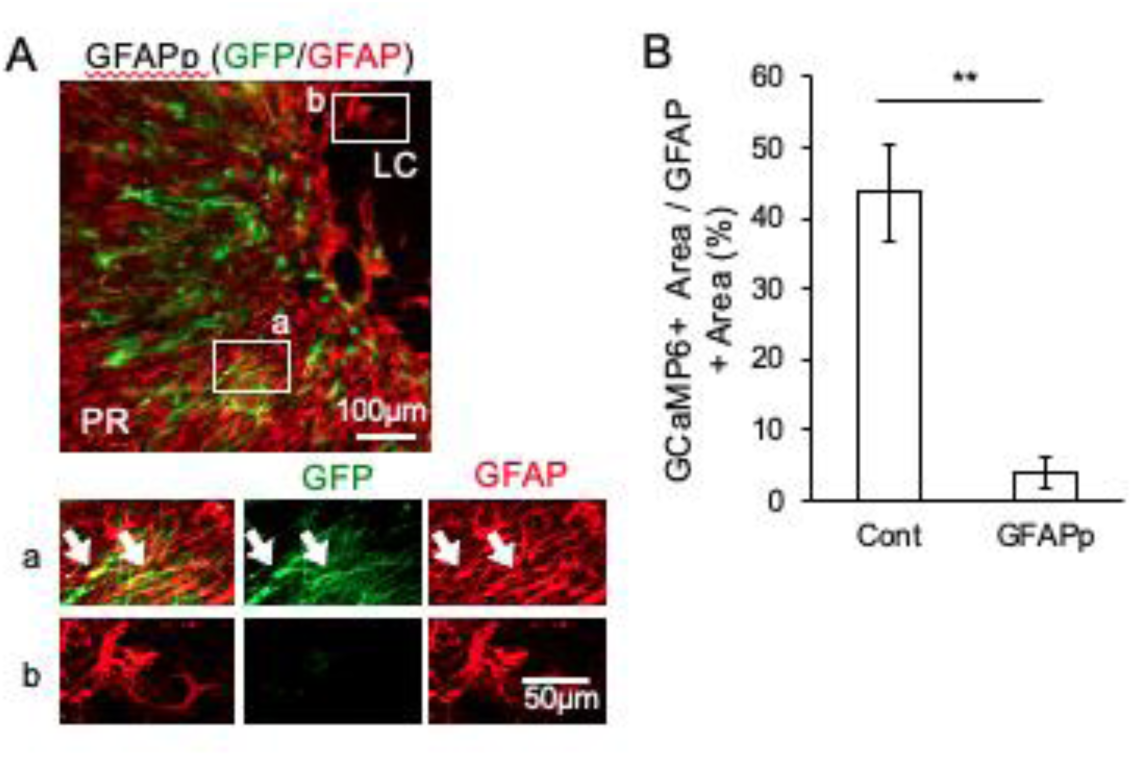
*Gfap* promoter labeling of a reactive astrocyte subpopulation distinct from scar-forming reactive astrocytes after brain injury. *Gfap*-CreERT2:ROSA26-GCaMP6 (GFAPp) mice were subjected to photo injury and administered tamoxifen at 1 dpi. A. Immunofluorescent images of GCaMP and GFAP at 14 dpi. Representative areas of the PR (a) and glial scar (b) highlighted by white boxes in the low magnification upper panel were enlarged in the lower panel. Arrows indicates GCaMP+/GFAP + cells in the PR. B. Proportions of areas occupied by cells labeled with GFP (GFP+ areas) in glial scars (GFAP+ areas). Mean ± SD, n = 3 or 4 mice, **P < 0.01 by Student’s t-test.

### 3.7 Fate of GFP+ cells

Because GFP+ cells are labeled by the *Nestin* promoter region, which is active in neural stem cells, they may undergo de-differentiation and possess multipotency in generating functional brain cells. To test this hypothesis, the expression of cell type markers was assessed in GFP+ cells of *Nestin-CreER^T2^: Rosa26-EGFP* mice in the PR at 28 dpi (Fig. 7). Most GFP+ cells were Nestin-, GFAP-and S100β-(Fig. 7A, B and C), whereas some were NG2+, PDGFRα+ and Olig2+ (Fig. 7D, E and F). A significantly higher proportion of GFP+ cells in the PR were positive for Olig2 (41 ± 6.2%) than for PDGFRα (10 ± 4.1%), NG2 (23.3 ± 2.9%), S100β (10 ± 0.0%), GFAP (1.25 ± 2.2%) and Nestin (1.6 ± 2.4%) (Fig. 7G). The lack of Nestin immunoreactivity indicated that GFP+ cells in the PR differed from GFP+ cells at 4 dpi. In addition, GFP+ cells were found to mature as oligodendrocytes, because significantly higher percentages were positive for Olig2 than for PDGFRα.

**Figure 7.**
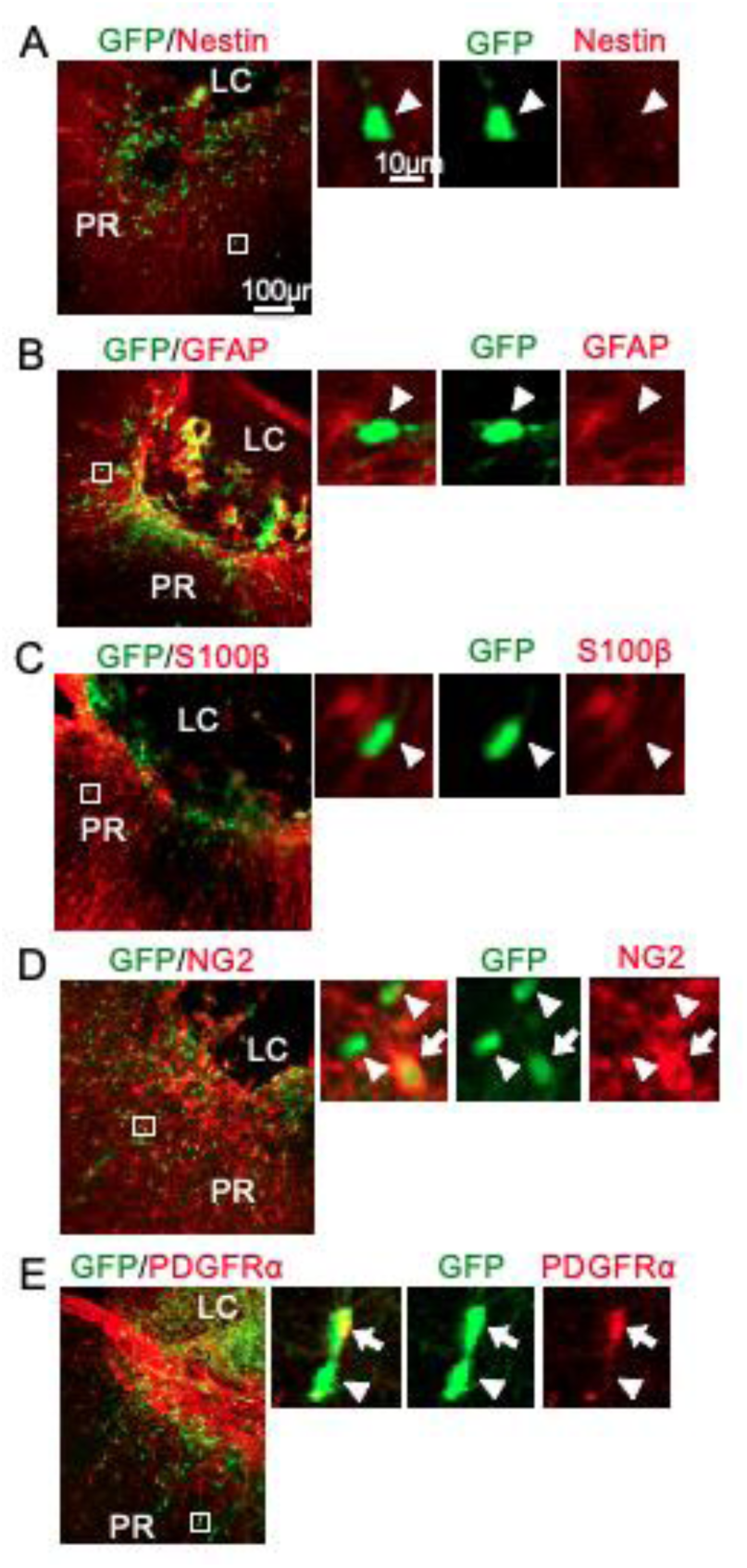
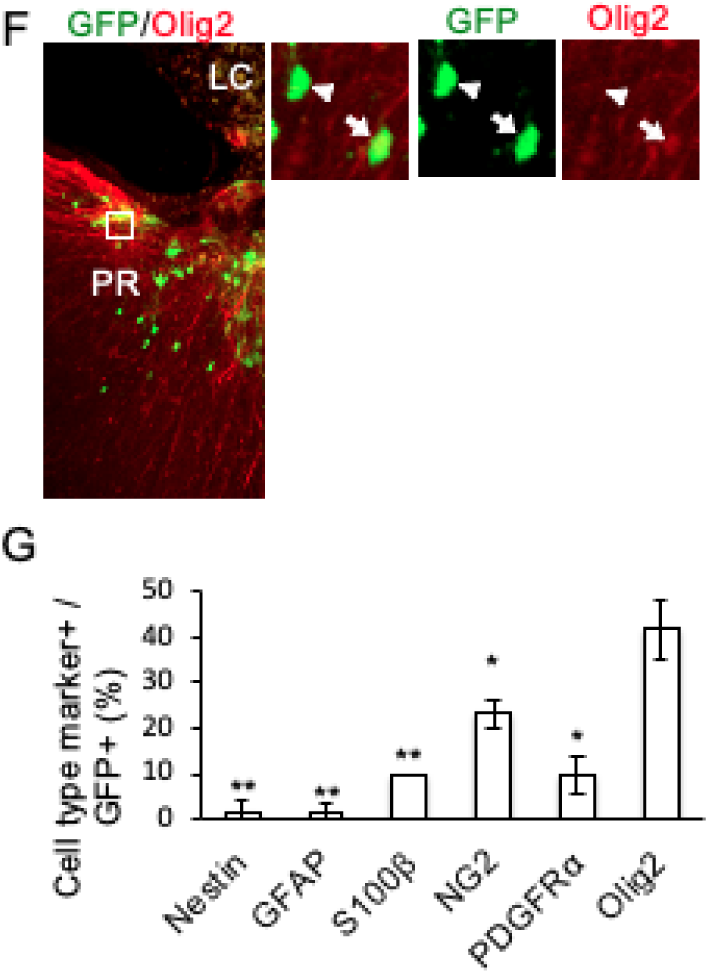
GFP+ cells in the PR mature as oligodendrocytes. *Nestin*-CreER^T2^:Rosa26-EGFP mice were subjected to photo injury and tamoxifen treatment at 1 dpi, followed by immunofluorescence analysis of GFP and cell type markers at 28 dpi. Representative images of GFP and Nestin (A), GFAP (B), S100β (C), NG2 (D), PDGFR (E) or Olig2 (F). Representative GFP+ cells highlighted by white boxes at low magnification images on the left-hand side are enlarged on the right hand side. G. Proportions of cell type marker positive cells in GFP+ cells. Mean ± SD, n = 60 GFP+ cells from three mice. *P < 0.05, **p<0.01 compared with Olig2 by ANOVA and Dunnett’s multiple comparison test.

## 4. Discussion

This study was initially designed to assess the origin, fate, and functions of Nestin-expressing reactive astrocytes by inducing photo injuries in the cerebral cortices of a mouse strain, in which tamoxifen-induced GFP expression was controlled by a *Nestin* promoter region commonly used to study neural stem/progenitor cells. We unexpectedly found, however, that OPCs were robustly labeled with GFP and were involved in glial scar formation. Glial scars are composed of reactive astrocytes derived from OPCs, but not from astrocytes, indicating that reactive astrocyte subtypes have distinct pathological implications. In addition, some GFP-labeled OPCs did not show upregulation of astrocyte markers and did localize in perilesional regions rather than glial scars. These cells were most likely committed to maturation as oligodendrocytes, but further studies are required.

The mouse postnatal brain contains NG2+ progenitor cells that produce both oligodendrocytes and astrocytes (Zhu et al., 2008), whereas NG2+ OPCs in normal adult brains produce oligodendrocytes alone (Kang et al., 2010; Nishiyama et al., 2009). Certain neurodegenerative disorders can lead to the differentiation of OPCs into reactive astrocytes, as observed in conditions such as spinal cord injury (Hackett et al., 2018) or stroke (Honsa et al., 2016; Kohyama et al., 2008). However, this differentiation is not consistent across all types of neural injuries, such as stab wound injuries (Komitova et al., 2011). Other studies investigating ischemic brain injury have noted a proliferation of NG2-expressing cells, identified as OPCs at the margin of the lesion (Janeckova et al., 2024). In our photo injury model, the *Nestin* promoter specifically labeled OPCs activated within the lesion core. This labeling allowed us to track the fate of activated OPCs, especially their differentiation into reactive astrocytes. This finding suggests a distinct fate determination pathway for OPCs in the lesion core compared to their behavior and role at the lesion periphery.

The *Nestin* promoter region, widely used to study neural stem cells, exclusively labeled OPCs after brain injury, despite Nestin immunoreactivity also being upregulated in reactive astrocytes and vascular cells (Suzuki et al., 2012). The expression of neural stem/progenitor cell markers, including Nestin, has been reported to be upregulated in astrocytes and pericytes after brain injury, with these cells becoming multipotent after isolation from lesions and maintenance in cell culture (Buffo et al., 2008; Nakata et al., 2017). Thus, they de-differentiate to multipotent cells after brain injury and subsequent exposure to a cell culture environment, explaining their rare activation of the *Nestin* promoter after brain injury alone. Alternatively, activation of the *Nestin* promoter in OPCs is consistent with their multipotency in producing reactive astrocytes.

The present study demonstrates that the *Nestin* promoter exclusively labeled an OPC subpopulation, which is activated by injury. Although the promoter regions of NG2 and PDGFRα have been used to label or manipulate OPCs (Honsa et al., 2016; Zawadzka et al., 2010), these genes are expressed in brain cells other than OPCs. For example, NG2 is expressed in pericytes and PDGFRα in perivascular and meningeal fibroblasts (Kelly et al., 2016; Stallcup, 2018). Only 50% of cells labeled by the NG2 promoter could be immunologically categorized as OPCs after brain injury (Hackett et al., 2016), suggesting that the NG2 promoter is active in non-OPC cells. *Nestin* promoter is also active in neural/progenitor stem cells in the subventricular zone (SVZ), and GFP-labeled cells have been found to migrate from the SVZ to the lesion site after photo injury (data not shown). However, we avoided the influence of these cells in quantitative analysis by excluding the region close to the SVZ. This highly selective activation of the *Nestin* promoter in OPCs after brain injury may be useful for the specific manipulation of pathologically-activated OPCs.

Some GFP-labeled OPCs in glial scars were converted to reactive astrocytes. The finding, that reactive astrocytes labeled with the *Gfap* promoter did not localize to glial scars, suggested that OPCs play a crucial role in glial scar formation. This hypothesis was supported by the impaired glial scar formation in STAT3 KO mice, which lack most GFP-labeled OPCs. Perilesional accumulations of proliferative reactive astrocytes are common after brain injuries; however the reactive astrocytes localizing at the interface between the LC and the PR have rarely been characterized. Because reactive astrocytes derived from OPCs are located exclusively at the interface between the LC and RP, they most likely insulate the LC physically from the PR and protect perilesional neurons, but are less likely to influence neurite growth and perilesional neuronal network regeneration. Conflicting findings have been observed in glial scars after spinal cord injury, in that neurites had to pass through the lesion site, including glial scars, for functional recovery.

Because glial scars after spinal cord injury interfere with neurite growth, the enzymatic reduction of the extracellular matrix produced by glial scars is a therapeutic strategy for improving outcomes (Morgenstern et al., 2002). Thus, the suppression of OPC activation and subsequent reduction of glial scar formation should enhance functional recovery after spinal cord injury, despite complicating the lesion itself due to its reduced physical insulation. The influence of glial scars on functional recovery after cerebral lesion must still be determined, mostly because long-term functional disability after photo injury has not been successfully measured to date.

GFP-labeled OPCs showed reduced expressions of Nestin and OPC markers by 28 dpi, although most of these cells continued to express Olig2. Thus, perilesional GFP-labeled cells, which did not have processes characteristic of myelinating oligodendrocyte during the period of investigation, are most likely in the process of oligodendrocytic maturation for remyelinating perilesional damaged of regenerated axons. Recent research using single-cell RNA sequencing on mice revealed that after focal cerebral ischemia, there is a significant reduction in mature myelinating oligodendrocytes by day 7 (Janeckova et al., 2024). This reduction gradually recovers over time, with the incidence of these cells approaching control levels by day 22. Based on our reserach, it was observed that OPCs within the lesion core differentiate into premyelinating oligodendrocytes in perilesional region, but do not contribute to the recovery of myelination. Thus, most of the newly myelinating cells are derived from OPCs located perilesional region.

Deletion of the *Stat3* gene reduced the number of GFP-labeled cells during early stages of injury, as well as reducing subsequent glial scars. STAT3 is essential for astrocyte activation (Ceyzériat et al., 2016), and the conditional knockout of STAT3 by *Nestin-Cre* was shown to impair the accumulation of Nestin positive reactive astrocytes after photo injury (Sawant et al., 2024). Thus, simultaneous deletion of STAT3 and labeling of GFP may reduce the numbers of GFP-labeled scar-forming reactive astrocytes without affecting perilesional GFP-labeled cells lacking astrocyte markers. However, STAT3 knockout reduced proliferating OPCs, even before the production of reactive astrocytes, robustly reducing the numbers of all GFP-labeled cells, both in the LC and PR. This reduced proliferation is consistent with results showing that the suppression of SOCS3, a suppressor of Jak-STAT3 signaling, after spinal cord injury upregulated the proliferation of OPCs (Hackett et al., 2016). STAT3 suppression may promote the death of OPCs after brain injury, because ciliary neurotrophic factor and leukemia inhibitory factor promote the survival of OPCs in cell culture via STAT3 (Steelman et al., 2016). This possibility could not be excluded, although few apoptotic cells detected by TUNEL staining were labeled with GFP regardless of STAT3 suppression (data not shown). The absence of cells doubly positive for TUNEL and GFP may reflect insufficient GFP labeling of apoptotic OPCs.

The current study demonstrates that glial scars were formed by reactive astrocytes derived from OPCs, rather than from astrocytes. The production of reactive astrocytes by OPCs required prior upregulation of Nestin. Thus, after brain injury, OPCs are converted to progenitor cells producing both oligodendrocytes and astrocytes, with similar progenitor cells, known as oligodendrocyte-type 2 astrocyte progenitors (O2A progenitor), obtained from developing brain (Barres et al., 1989; Nishiyama et al., 1996). Thus, the results of the present study indicate that OPCs likely converted to O2A progenitors after brain injury. Further investigation on the cell signaling pathways underlying this conversion may lead to more accurate diagnosis and better treatment of brain injury.

## CRediT authorship contribution statement

**Hinata Matsuda:** Investigation, Writing – original draft. **Erika Tsuji**: Investigation. **Agnes A. O. Riberu**: Writing – Review & editing. **Hideyuki Okano**: Resources, Writing – Review & editing. **Mitsuhiro Morita**: Supervision.

## Data availability

Data will be made available on request.

## Acknowledgement

We are grateful to Drs. R. Kageyama, J. Miyazaki and F. Kirchhoff for generous supply of *Nestin-CreER^T2^*, *Rosa26-GFP* and *Gfap-CreER^T2^* mice respectively, and Dr. S. Miyata for providing Guinea pig anti-GFAP antibody. We also express our appreciation to Dr. K. Inoue and Mr. D. Hatano for scientific discussion and manuscript preparation, respectively.

